# Transcriptomics effects of per- and polyfluorinated alkyl substances in differentiated neuronal cells

**DOI:** 10.1101/2024.01.26.577438

**Authors:** Logan Running, Judith R. Cristobal, Charikleia Karageorgiou, Michelle Camdzic, John Michael Aguilar, Omer Gokcumen, Diana S. Aga, G. Ekin Atilla-Gokcumen

## Abstract

Per- and polyfluorinated alkyl substances (PFAS) are pervasive environmental contaminants that bioaccumulate in tissues and pose risks to human health. Increasing evidence links PFAS to neurodegenerative and behavioral disorders, yet the underlying mechanisms of their effects on neuronal function remain largely unexplored. In this study, we utilized SH-SY5Y neuroblastoma cells, differentiated into neuronal-like cells, to investigate the impact of six PFAS compounds— perfluorooctanoic acid (PFOA), perfluorooctanesulfonic acid (PFOS), perfluorodecanoic acid (PFDA), perfluorodecanesulfonic acid (PFDS), 8:2 fluorotelomer sulfonate (8:2 FTS), and 8:2 fluorotelomer alcohol (8:2 FTOH)—on neuronal health. Following a 30 μM exposure for 24 hours, PFAS accumulation ranged from 100–7500 ng/mg of protein. Transcriptomic analysis revealed 721 differentially expressed genes (DEGs) across treatments (p_adj_ < 0.05), with 11 DEGs shared among all PFAS exposures, indicating potential biomarkers for neuronal PFAS toxicity. PFOA-treated cells showed downregulation of genes involved in synaptic growth and neural function, while PFOS, PFDS, 8:2 FTS, and 8:2 FTOH exposures resulted in upregulation of genes related to hypoxia response and amino acid metabolism. Lipidomic profiling further demonstrated significant fatty acid upregulation with PFDA, PFDS, and 8:2 FTS, alongside triacylglycerol downregulation with 8:2 FTOH. These findings suggest that the neurotoxic effects of PFAS are structurally dependent, offering insights into the molecular processes that may drive PFAS-induced neuronal dysfunction.

**Synopsis:** Per- and poly fluorinated alkyl substances (PFAS) have been shown to bioaccumulate in human tissues and affect health. This study aims to provide insights into the specific biological processes through which PFAS exposure affects neuronal cells.

## INTRODUCTION

Per- and polyfluorinated alkyl substances (PFAS) are a class of chemical contaminants that have attracted global concern due to their persistence in the environment, widespread use, and potential adverse effects on human health. Although monitoring efforts over the past few decades have led to positive trends, such as the phase-out of legacy compounds like perfluorooctanoic acid (PFOA) and perfluorooctanesulfonic acid (PFOS)^1,2^, the production and use of PFAS remain significant. ^3^ The US Environmental Protection Agency (EPA) reports the use of thousands of PFAS with various structures differencing at the head group, fluorination state, and branching. ^4^ Recent studies from the United States show that detectable levels of PFAS are found in the serum of 97% of the population, demonstrating their ubiquitous environmental presence and highlighting ongoing human exposure risks.^5^

PFAS are highly bioaccumulative due to their inherent chemical stability^6,7^ and fat solubility^8^, resulting in prolonged serum half-lives in humans (ranging from months to over five years depending on the PFAS type)^9,10^. PFAS exposure routes include ingestion, inhalation, and skin contact, with certain professions (e.g., firefighters, industrial workers, and ski board waxers) being at heightened risk.^11^ The bioaccumulation of PFAS has been observed in multiple organs, including the brain, liver, and serum.^12^ Notably, PFAS can cross the blood-brain barrier (BBB) and accumulate in brain tissue^13,14^, posing potential neurotoxic risks.

PFAS are known to disrupt several biological processes, including endocrine function^15, 16^, reproductive health^17^, and lipid metabolism^18,19^. Bioaccumulation and hepatotoxicity is a well-documented outcome of PFAS exposure, with studies linking these compounds to liver dysfunction^20,21^, and certain cancers, such as liver and testicular cancers, have been associated with PFAS exposure^22,23^.

PFAS are detected in brain tissue and cerebrospinal fluid.^24,25^ Perfluorocarboxylates and perfluorosulfonates with 6-14 carbon chains were detected in various organisms, with the longer chain PFAS generally occurring at higher concentrations.^14^ Further, PFHxA and PFBA have been detected in brain up to 300 ng/g in humans^26^. A recent study found PFOA, perfluorohexane sulfonic acid (PFHxS), and PFHxA in various parts of the male brain, with levels correlating strongly with PFAS levels in respective blood samples from these individuals.^27^ Surprisingly, the same study showed that low levels of PFOS in the blood and not in the brain, indicating selective accumulation of different PFAS in the brain.^27^

Previous studies have suggested that PFAS accumulation in the brain could be mediated by lipid transporters^14^ other organic anionic transporter,^28^ and is linked to the disruption of the BBB^29^. Studies have linked PFAS neurotoxicity to changes in neurotransmitter levels and disruption of synaptic homeostasis, which in turn, can serve as risk factors for neurodegenerative diseases, such as Alzheimer’s or Parkinson’s disease.^30,31^ Further, infant exposure of PFAS during pregnancy and childhood through the mother’s blood and breast milk have been linked to dysregulations in childhood physiology and behavior.^32,33^ PFOS, PFHxS, and perfluorononanoic acid (PFNA) are linked to several symptoms of attention deficit hyperactivity disorder (ADHD), such as hyperactivity and aggression. In contrast, PFHxS was correlated with internalizing behaviors such as depression and anxiety.^34^ Despite these findings, the molecular mechanisms underlying PFAS-induced neurotoxicity remain poorly understood.

In this study, we aimed to investigate the neurotoxic effects of six long-chain PFAS (PFOA, PFOS, perfluorodecanoic acid [PFDA], perfluorodecanesulfonic acid [PFDS], 8:2 fluorotelomer sulfonate [8:2 FTS], and 8:2 fluorotelomer alcohol [8:2 FTOH]) using differentiated SH-SY5Y neuroblastoma cells. These cells were exposed to PFAS at sub-toxic concentrations for 24 hours to simulate levels observed *in vivo*. By analyzing changes in gene expression and lipid levels, we sought to identify the specific biological processes affected by PFAS exposure. Our results demonstrate structure-dependent differences in PFAS uptake and toxicity, with distinct profiles of transcriptomic changes observed for each compound. Lipidomic analysis also suggested differences in the changes in lipid levels based on PFAS’ structure. These findings provide new insights into the molecular mechanisms of PFAS neurotoxicity and their potential impact on neuronal health.

## MATERIALS AND METHODS

### Materials

PFOA, PFOS, PFDA, PFDS, 8:2 FTOH, and 8:2 FTS were purchased from Millipore Sigma^®^ (Burlington, MA). Standard for 8:2 FTOH was obtained from Toronto Research Chemicals (Toronto, ON, Canada). ^13^C isotopically labeled standards including M8PFOA, M6PFDA, M8PFOS, M2-8:2 FTS, M4PFOS, ^2^H_2_-^13^C_2_-6:2 FTOH and ^13^C_2_-6:2 FTOH were purchased from Wellington Laboratories (Guelph, ON). Dansyl chloride (>99%), 4-(dimethylamino)pyridine (>99%), and N,N-diisopropylethylamine (>99%) were purchased from Sigma-Aldrich (Saint Louis, MO). ACS-grade ammonium formate was purchased from Alfa Aesar (Ward Hill, MA). Dichloromethane, hexane, and formic acid (88%) were purchased from Fisher Scientific (Waltham, MA). Ammonium acetate, LC-MS grade methanol and HPLC grade methanol were purchased from J.T. Baker^®^ (Radnor Township, PA). Type I water (18.2 MΩ-cm) was generated using using a Barnstead Nanopure^TM^ Diamond (Waltham, MA) purification system.

SH-SY5Y neuroblastoma cells were purchased from the American Type Culture collection (Manassas, VA). Dulbecco’s modified Eagle media (DMEM) / Hams-F12 50:50, fetal bovine serum (FBS), trypsin, and penicillin/streptomycin antibiotic cocktail were purchased from Corning^®^ (Corning, NY). GlutaMAX^TM^ and Neurobasal^TM^ media were purchased from Gibco (Billings, MT). Thiazolyl Blue Tetrazolium Bromide was purchased from VWR (Radnor, PA). B-27 100X supplement was purchased from Fisher Scientific (Waltham, MA). A lactate dehydrogenase (LDH) chemical assay kit, and all-trans retinoic acid were purchased from Cayman Chemical Company (Ann Arbor, MI). Potassium Chloride was purchased from Mallinckrodt (Staines-upon-Thames, United Kingdom). Brain derived neurotrophic factor (BDNF) was purchased from Prospec^TM^ Bio (Rehovot, Israel).

## Methods

### Cell culture

SH-SY5Y cells were cultured based on instructions provided by American Type Culture Collection. Cells were grown in 10 cm plates and allowed to reach a confluency of 85% after approximately 5-6 days. After reaching 85% confluency cells were sub-cultured to new 10 cm plates at a 1:10 ratio. PBS was used to rinse cells before sub culturing and trypsin was used to uplift cells from plate. Cellular media was replaced every 3-4 days as needed.

### Cellular differentiation

Cells were differentiated over an 11-day period using retinoic acid and brain derived neurotrophic factor.^35^ Over the 11 days, three different media compositions are used. The first was the base media used for general cell culturing for SH-SY5Y cells composed of DMEM/F12 50:50 media with 10% FBS and 1% penicillin/streptomycin (P/S). The second is stage 1 media which used DMEM/F12 50:50 as the base supplemented with 2.5% FBS, 1% P/S, and 10 μM all-trans retinoic acid. Lastly, stage 2 media which used neurobasal media supplemented with 1X B-27, 20 mM KCl, 1% P/S, 2 mM glutaMAX, and 50 ng/mL BDNF. Differentiation over 11 days was carried out as described in the literature.^35,36^

### PFAS exposure and cell collection

After the 11-day differentiation SH-SY5Y cells in 10 cm plates were exposed to 30 μM PFAS PFOA, PFOS, PFDA, PFDS, 8:2 FTOH, and 8:2 FTS at a final concentration of 1% MeOH. Differentiated SH-SY5Y cells were also treated with only 1% MeOH as a vehicle control. Treatments lasted 24 hours and were done in triplicate. Cells were washed with cold 1x PBS and collected by centrifugation at 4°C, 300 rcf, 5 minutes. Cell pellets were resuspended with 500 μL of cold 1x PBS. An aliquot of 50 μL was set aside for protein normalization and the rest of the cell pellets were collected and stored for PFAS extraction and analysis.

### Viability assays

After day 11, treatments of PFAS in MeOH were added for final concentrations of 3, 10, 30, and 100 μM with 1% MeOH with a vehicle control of 1% MeOH. An additional treatment 10% Triton^TM^ was added as a positive control for membrane permeabilization (n=5). Treatments lasted 24 hours.

MTT (3-(4,5-dimethylthiazol-2-yl)-2,5-diphenyltetrazolium bromide) assays were also performed. After a 24-hour PFAS treatment, cell plates were centrifuged at 200xg for 2 minutes at 24°C. Media was removed and 200 μL of 9% MTT in base media was added back. The plates were incubated at 37°C under 5% CO_2_ for 3 hours. After the plate was centrifuged a 150 μL portion of media was removed and 90 μL of DMSO was added back. Plates were then incubated at 37°C for 15 minutes and then mixed gently to dissolve formazan crystals. Absorbance was measured at 550 nm in a plate reader.

### PFAS extraction and analysis

PFOA, PFOS, PFDA, PDFS and 8:2 FTS extraction and analysis were carried out based a modified protocol described from a previously published protocol.^36^ Cell pellets treated with PFAS along with the vehicle control were resuspended in 1 mL cold MeOH spiked with corresponding ^13^C-mass-labelled isotopically labelled standards to a final concentration of 100 ppb. The cell pellets were vortexed for 30s and sonicated at 40% power three times for 10s while on a cold metal block. The samples were centrifuged (16,000g, 10 min, 4°C). 900 μL of the supernatant was transferred to a 1 mL dram glass vial without disrupting the cell pellet. The extraction procedure was done twice. The combined supernatant (1800 μL) was dried down under a nitrogen stream. The samples were reconstituted in MeOH spiked with 100 ppb each of M4PFOA and M4PFOS as internal standards. The volume of MeOH for reconstitution was based on protein content to ensure equal loading.

Cell pellets treated with 8:2 FTOH were extracted with 1mL cold acetonitrile instead of MeOH. The extraction was done similarly with other PFAS-treated pellets. However, the extraction was only done once and M2-8:2 FTOH was used as standard. The 8:2 FTOH acetonitrile extract (900 μL) was not dried down and was derivatized using dansyl chloride as described in ref^37^ and below.

### FTOH derivatization

After extraction of 8:2 FTOH with acetonitrile, 900 μL of the extract was transferred to a 1.5 mL amber vial and mixed with 200 μL of 10 mg/mL dimethylaminopyridine (DMAP) and 10 mg/mL dansyl chloride in dichloromethane (DCM) with 1% N,N-diisopropylethylamine. The reaction mixture was vortexed for 30s and incubated at 65°C for 1 hour. Following this, incubation reactions were quenched on ice for 5 min. The reaction mixtures were transferred to 15 mL falcon tubes and combined with 3 mL of water and extracted with 5 mL hexane. The mixture was vortexed then centrifuged at 4200xg for 15 min at 4°C. The top layer was set aside and a second extraction was done with an additional 5 mL of hexane. Approximately 10 mL of combined extracts were further extracted by solid phase extraction on a silica cartridge. The Waters Silica cartridge was equilibrated with 4 mL DCM and 4 mL hexane. The combined extracts were loaded into the pre-conditioned cartridge and dried under vacuum for 5 min. The cartridge was eluted with 8 mL of 1:1 hexane/DCM without vacuum. The eluent was collected in a 15 mL falcon tube and dried under nitrogen stream. The samples were resuspended with 200 μL of starting mobile phase and analyzed via Triple Quadrupole LC-MS/MS.

### LC-QTOF-MS analysis

PFOA, PFOS, PFDA, PFDS, and 8:2 FTS analyses were done by using liquid Chromatography-quadrupole time of flight mass spectrometry (LC-QToF-MS). Separation was done using a Gemini C18 reversed-phase column (5 μM, 4.6 mm x 50 mm, Phenomenex). Analyses were carried out in negative mode using an Agilent 1260 HPLC in tandem with an Agilent 6530 Jet Stream ESI-QToF-MS system with a Gemini C18 reversed-phase column (5 μM, 4.6 mm x 50 mm, Phenomenex). Mobile Phase A was composed of 5 mM ammonium acetate in water at pH 3.80 while mobile phase B was 100% MeOH. The gradient for PFAS elution and separation began with 50% B at 0.5 mL/min and after 5 min increasing to 95% until 15 min. The gradient was held at 95% B until 30 min and switched back to 50% B until 35 min for equilibration. Injection volume is 5 μL and flow rate is 0.5 mL/min. The capillary voltage and fragmentor voltage were set to 3500 V and 175 V, respectively. The drying gas temperature was set to 350°C with the flow rate set to 12 L/min. All PFAS were detected in negative mode as [M-H]– except for PFOA, PFDA, M4PFOA, M8PFOA, and M6PFDA at [M-COOH]–^38^. A list of *m/z* adducts can be found in Table S1.

Quantification of PFAS was done by isotope dilution. Prior to extraction, the corresponding ^13^C mass-labelled isotope standard (surrogate) of each PFAS analyte was spiked on the cell pellets resuspended in methanol or acetonitrile. The final concentration of the surrogate standard is expected to be 90 ppb after extraction and reconstitution of PFAS extracts with methanol spiked with 100 ppb M4PFOA and 100 ppb M4PFOS as internal standards.

Equation 1 below shows the equation used for isotope dilution.

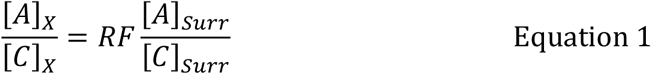

where: *RF* = 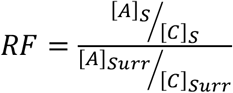

*Equation 1: Isotope dilution. X=unknown, S=Standard, A=Peak Area, C=Concentration, RF= Response Factor*

### Liquid Chromatography-Tandem Mass Spectrometry (LC-MS/MS)

Derivatized FTOH quantification was performed using an Agilent 1200 HPLC coupled to a Thermo Scientific^TM^ TSQ Quantum Ultra triple quadrupole mass spectrometer (LC-MS/MS) operated in the positive mode electrospray ionization (+ESI) with a spray voltage of +3000 V, vaporizer and capillary temperature of 300 °C. Nitrogen was used as the sheath gas (35 arb.) and auxiliary gas (30 arb.). Precursor ions, product ions and collision energies are summarized in Table S1. Separation was achieved using a Restek Raptor C_18_ analytical column (2.7 μm particle size, 100 x 3 mm) (Restek, Bellefonte, PA), and an aqueous mobile phase (A) of 5 mM ammonium formate with 0.1% formic acid in Nanopure^TM^ water, an organic mobile phase of 5 mM ammonium formate with 0.1% formic acid in methanol (B), and a flow rate of 0.200 mL/min. A 40-minute gradient method was used to separate and analyze the derivatized FTOH starting with 55% B for 3 minutes, ramping to 95% B over 13 minutes and held at 95% B for 18 minutes. The mobile phase was then returned to starting conditions of 55% (B) over 1 minute and held for 7 minutes for equilibration before subsequent injection. The analyte, mass-labelled standards, transitions, and collision energies are shown in Table S2. Quantification was performed using the isotopic dilution. The final concentration of the surrogate standard is expected to be 50 ppb after extraction and reconstitution of PFAS extracts with starting mobile phase.

### Combustion Ion Chromatography (CIC) Analysis and Quantification of 8:2 FTOH in neurobasal media

A 100 μL of MeOH as control and 3 mM stock 8:2 FTOH were spiked in 10 mL neurobasal media supplemented with 50 ng/mL BDNF, 2 mM Glutamax, 20 mM KCl, 1x B-27, and 1% P/S contained in 10 cm dishes. The samples were incubated for 24 hrs at 37°C and 5% CO_2_. A 400 μL aliquot of each sample (n=3) was obtained at various time points from 0 to 24 hrs. The aliquots (200 μL) were passed through an adsorbable organohalide (AOX) system and washed with 5000 ppm KNO_3_ to remove any other organic and inorganic contaminants leaving only organofluorine. The carbon material in the AOX was transferred to CIC sampling boats and analyzed via the CIC instrument. Fluoride concentration of 8:2 FTOH extracts were determined using CIC instrument equipped with AQF-2100H combustion system coupled with a ThermoScientific Dionex ICS-6000 DC ion chromatography system. Argon and oxygen were used as a carrier gas and combustion gas, respectively. The flowrates for argon was set to 200 mL/min while oxygen was set to 400 mL/min. Potassium hydroxide (KOH) was used as the mobile phase. Anions separation were achieved using a Dionex IonPac^TM^ AS18-Fast column (2 x 150 mm, 4 μM) and 0.25 mL/min flowrate and varying concentrations of KOH for the gradient program. The ion chromatography gradient elution program started with 10 mM KOH held for 4 mins, then increased to 40 mM and held for 3 minutes. It was switched back to 10 mM KOH for 1 min. The total run time is 13 mins with equilibration time for 4 mins. The quantification of percent remaining 8:2 FTOH in the media was calculated based on an external calibration curve.

### RNA extraction, library preparation and sequencing

Frozen cell pellets were transported on dry ice to the University at Buffalo Genomics core and stored at -80°C until analyzed. RNA was extracted using TriZol using Invitrogen protocols. Quality of RNA was assessed using Agilent’s Fragment Analyzer and Qubit. RNA was then used to prepare an RNA library using an Illumina RiboZero Total Stranded RNA library prep kit. Libraries were pooled together at 10 nM with concentration being determined with a QuantaBio Universal qPCR reaction kit. The pooled library was sequenced using a NovaSeq6000 (PE100).

### RNA-seq analysis

The obtained RNA sequences were evaluated using FastQC^39^ and MultiQC^40^ for quality control. Two samples were eliminated from the downstream analysis due to evidence of contamination. For five out of the six treatments, we have at least three samples, a sample was discarded each from the control and the FTOH treatment category, leaving two samples for the downstream analysis for each category. The two discarded samples were flagged as contamination due to their low number of successfully assigned alignments: for Control_3_S29 less than 14.2% of the reads were successfully mapped to the reference, while for FTOH_3_S47 a mere 14% of the total number of reads were successfully aligned. FastQC screen reports for the two samples confirmed that there are discrepancies with these samples either due to contamination or issues with the library preparation due to the presence of reads that produce multiple hits to multiple genomes. The RNA-seq reads were mapped to the human transcriptome reference (hg38) from Ensemble using hisat2^41^ and quantified using featureCounts^42^. The differential expression analysis was carried out using DESeq2.^43^ DESeq2 estimates the log fold-change of transcription for each gene using the Wald test of significance and performs multiple hypotheses correction on the raw read counts. DESeq2 can estimate dispersion with one ‘degree of freedom,’ calculated as the difference between the number of samples and the number of coefficients. In the context of our study after the removal of the two contaminants, there are four samples and two groups (represented by two coefficients), thus, two ‘degrees of freedom’ are available. However, it is important to note that statistical power diminishes rapidly in this context, leading to a decreased sensitivity, thereby hindering the detection of smaller log-fold changes.^44^ To determine the significantly differentially expressed genes we filtered our results for an p_adj_ of 0.05 (where the p_adj_ we refer to the multiple-hypotheses-corrected p-value).

The fastq RNA-seq data have been submitted to GEO and the project name will be provided once available.

### Lipid extraction and analysis

Lipid extraction and sample preparation were carried out as we described previously.^45, 46^ Samples were normalized based on their protein content and resuspended in spiked chloroform. C17:0 sphingomyelin, C^13^ oleic acid, C17:0 ceramide, C57:0 TAG, and C18:0/C18:0 phosphatidylcholine-d^70^ were used as internal standards. LC-MS-based lipid analysis analyses were carried out using an Agilent 1260 HPLC in tandem with an Agilent 6530 Jet Stream ESI-QToF-MS system as we described previously. Lipid species were targeted by extracting the corresponding *m/z* for each ion in MassHunter Qualitative Analysis software (v11). Peak areas for each ion were integrated and represented as abundance. Relative abundances used in heatmaps for the PFAS treatments (n=3) were normalized to the vehicle control by dividing the average abundance of a lipid species in the vehicle control by the individual abundances of each PFAS treatment sample.

## RESULTS AND DISCUSSIONS

### Effect of PFAS exposure on cell viability of differentiated SH-SY5Y cells

The SH-SY5Y cells were differentiated as we described previously^36^ and then treated with PFOA, PFOS, PFDA, PFDS, 8:2 FTOH, and 8:2 FTS for 24 hours at concentrations of 3, 10, 30, and 100 μM. Cell viability was assessed using MTT assay. None of the treatments significantly reduced cell viability (p > 0.05), except for 8:2 FTS, which induced a modest cell death at 100 μM (p < 0.05, approximately 30%, **Figure 1**).

**Figure 1:**
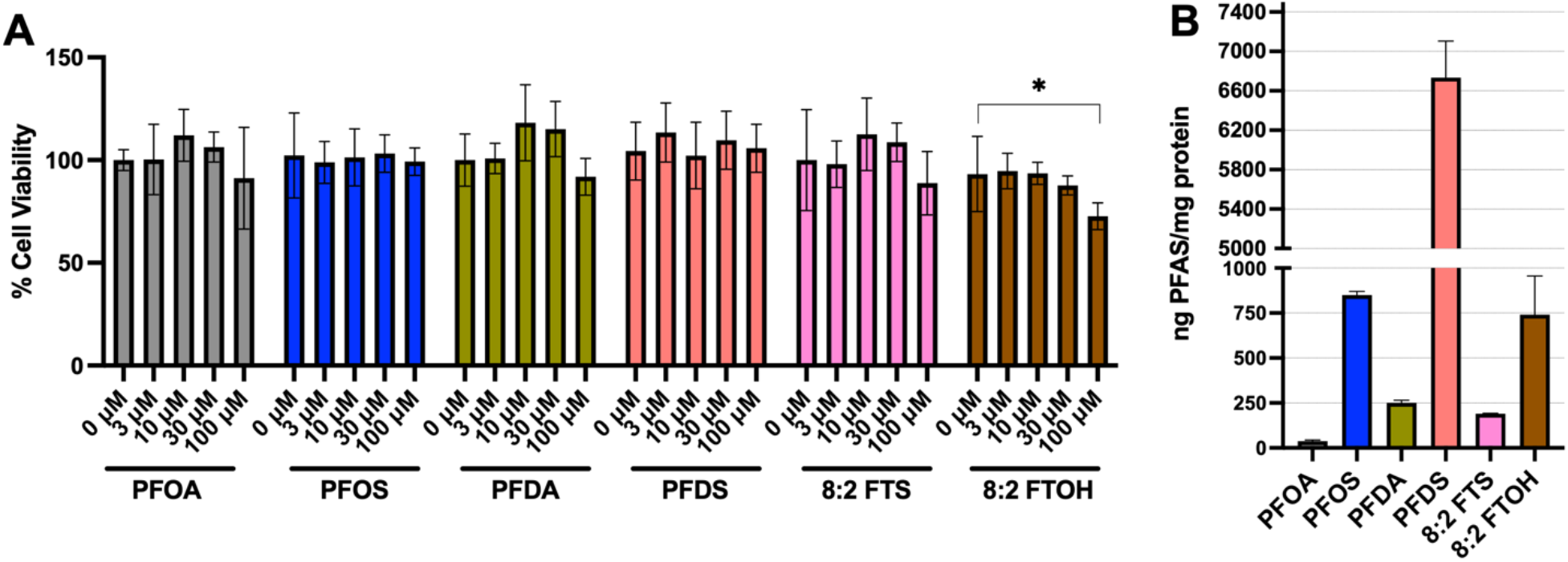
(**A**) Viability of SH-SY5Y cells treated with 3, 10, 30, and 100 μM PFOA, PFOS, PFDA, PFDS, 8:2 FTOH, and 8:2 FTS for 24 hrs. 8:2 FTOH treatment at 100 μM showed a slight decrease in viability, which was statistically significant based on a paired t-test (* p < 0.05). (**B**) Uptake of PFAS in differentiated SH-SY5Y cells after a 30 μM exposure for 24 hrs (n=3). The error bars represent % standard deviation.

One challenge in PFAS exposure studies is replicating environmental PFAS exposure conditions in the laboratory, as PFAS exposure is typically chronic in nature. ^19^ PFAS are frequently detected in human serum at low part-per-billion (ppb) or nanomolar concentrations^47, 48^, but do not exhibit cytotoxicity in cultured cells at concentrations up to 100 μM^15, 49, 50^, consistent with our findings. Additionally, PFAS are known to reversibly bind to proteins, which can decrease their bioavailability for uptake *in vitro*.^51^ Based on our viability results, we selected 30 μM as the treatment concentration for subsequent experiments, as it was the highest concentration that did not induce significant toxicity.

### PFAS uptake in differentiated SH-SY5Y cells

PFAS uptake in differentiated SH-SY5Y cells treated with 30 μM PFAS was determined via isotope dilution method (see methods section for details). Briefly, cell pellets were collected, and protein contents were measured, followed by LC-MS analysis. **Figure 2** shows the average uptake of PFOA, PFOS, PFDA, PFDS, 8:2 FTS and 8:2 FTOH. Results show that the cellular uptake of PFAS tested increases with increasing chain length for both the carboxylate and sulfonate head groups. Specifically, PFDA and PFDS (10 carbon chain length) are taken up at higher levels than PFOA and PFOS (8 carbon chain length). Moreover, the fully fluorinated sulfonate, PFDS, is taken up more effectively than its fluorotelomer counterpart, 8:2 FTS.

**Figure 2:**
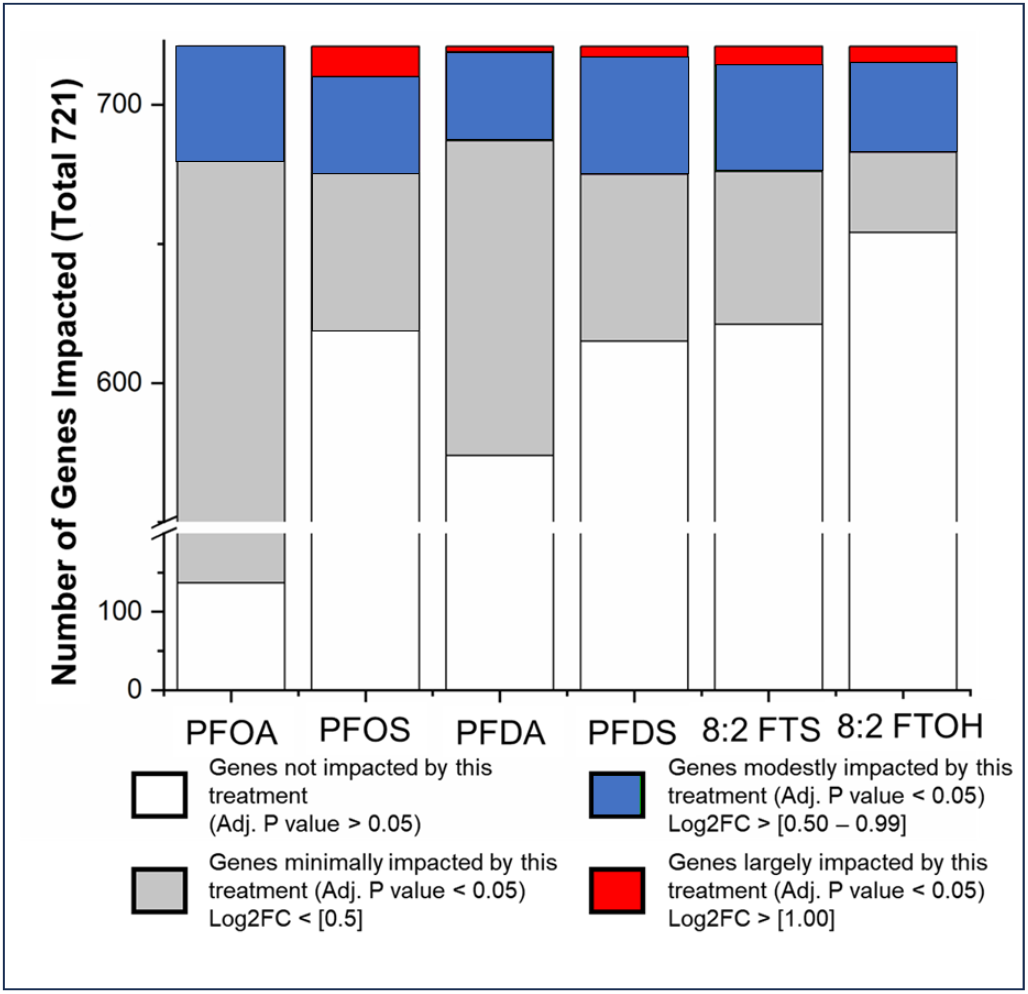
Distribution of log_2_FC observed for gene expression in SH-SY5Y cells after exposure to six different PFAS species. Log_2_FC > [0.5] were selected. Log_2_FC of > [1] was chosen to represent genes that were doubled or halved in expression and comprised less than 1% of the genes impacted by the PFAS treatments.

Studies have shown that fluorotelomer alcohols (FTOHs) have higher volatility compared to carboxylate and sulfonate-containing PFAS and can be transformed into other PFAS, primarily their corresponding carboxylate counterparts.^52^ In our study, PFOA was not detected in 8:2 FTOH-treated cells, indicating that 8:2 FTOH was not metabolized under treatment conditions. However, we observed a considerable decrease in 8:2 FTOH concentration in the treatment media over time, with ∼10% remaining in solution after 24 hours of exposure, respectively (see methods section for details). This decrease might indicate potential evaporation of 8:2 FTOH during exposure, making direct comparisons of uptake between 8:2 FTOH and other PFAS complicated . Nevertheless, we found that 8:2 FTOH accumulated in differentiated SH-SY5Y cells at levels much higher than 8:2 FTS and comparable to PFOS. These findings suggest that both chain length and headgroup composition significantly influence PFAS uptake in differentiated SH-SY5Y cells. Among the PFAS tested, uptake increases with chain length, with fully fluorinated sulfonates showing the highest cellular uptake, while short-chain carboxylates exhibited reduced uptake.

### Changes in gene expression in differentiated SH-SY5Y cells

To assess the impact of PFAS exposure on neuronal cells, we performed a transcriptomic analysis to identify pathways that were differentially regulated following PFAS treatment. Differentiated SH-SY5Y cells were treated with 30 μM PFOA, PFOS, PFDA, PFDS, 8:2 FTOH, and 8:2 FTS. Cells were harvested as detailed in the methods section. After mRNA extraction (n=3 per condition), the samples were sequenced using Illumina sequencing technology (see methods section for details). Gene expression levels in treated cells were compared to controls and expressed as log_2_ fold change (Log_2_FC). Genes showing significant differential expression (p_adj_ < 0.05) were compiled into a list for further statistical analysis and Gene Ontology (GO) enrichment analysis.

**Figure 2** illustrates the total pool of differentially expressed genes impacted by all six PFAS treatments compared to the control (p_adj_ < 0.05), as well as the distribution of genes affected per treatment. Each bar represents an individual treatment, with colors indicating the magnitude of gene expression changes. Gray represents genes with a log_2_ fold change (log_2_FC) < 0.5, blue represents genes with log_2_FC between 0.5 and 0.99, and red represents genes with log_2_FC > 1. White sections indicate genes that were not significantly affected by the corresponding treatment (p_adj_ > 0.05).

In total, 721 unique genes were impacted by the combined PFAS treatments. Among the treatments, PFOA affected the largest number of genes, significantly altering the expression of 584 genes (p_adj_ < 0.05). In contrast, the remaining five PFAS affected approximately 100 genes each (p_adj_ < 0.05): PFOS (102 genes), PFDS (106 genes), PFDA (147 genes), 8:2 FTS (100 genes), and 8:2 FTOH (67 genes). Interestingly, despite PFOA showing the lowest cellular uptake compared to the other PFAS (**Figure 1**), it resulted in the greatest number of differentially expressed genes, although the magnitude of these changes was relatively small (log_2_FC < 0.5). Conversely, PFOS, which exhibited higher cellular uptake, affected fewer genes (**Figure 2**). A full list of genes separated by treatment group and sorted by log_2_FC is provided in **Table S3**.

To visualize the overlap of differentially expressed genes between the treatment groups, an upset plot was generated (**Figure 3A**). This plot groups genes based on the number of treatments in which they are commonly affected. Of the 721 differentially expressed genes, 425 were exclusive to PFOA treatment. Other PFAS treatments uniquely affected between 13 and 36 genes, while 193 genes were differentially expressed across combinations of PFAS treatments. Notably, PFDA and PFOA affected the expression of 53 common genes, which may be attributed to the shared carboxylate headgroup. Additionally, 11 genes (10 protein-coding and one RNA-coding) were differentially regulated across all six treatments (**Figure 3B**).

**Figure 3:**
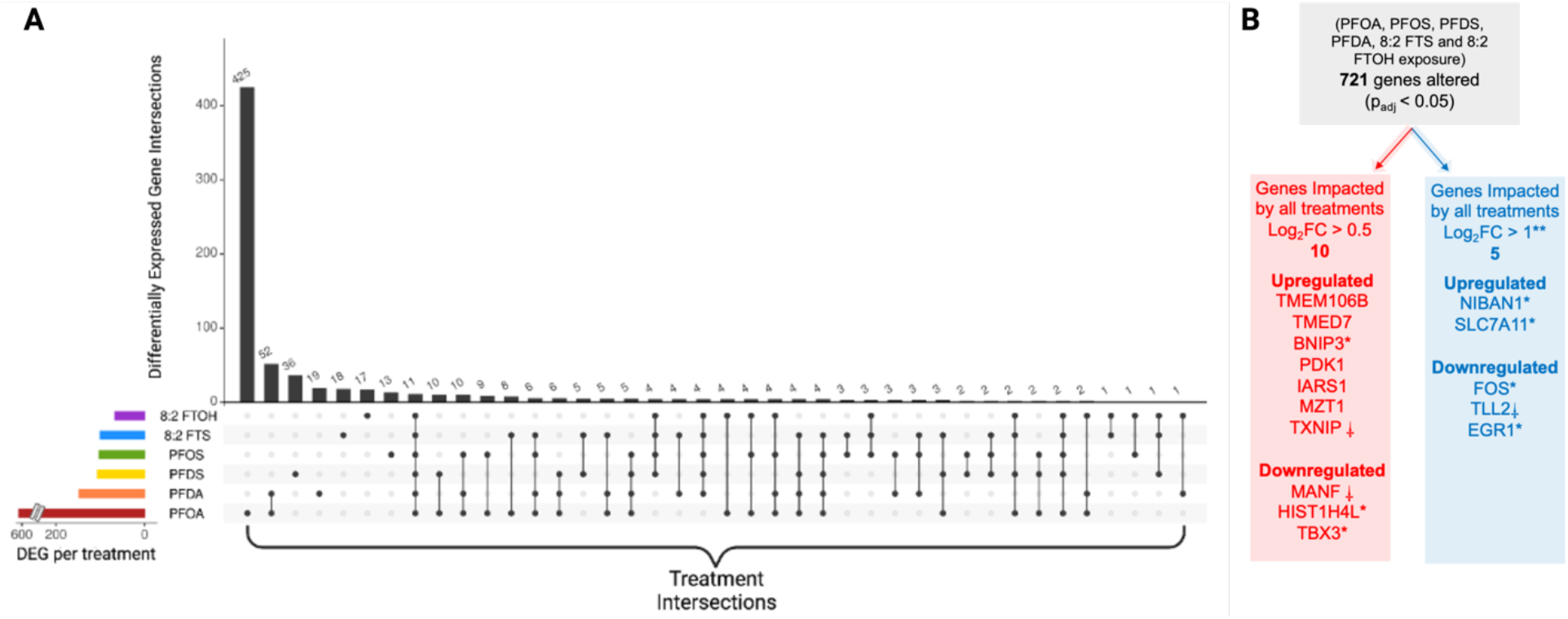
**A**. An upset plot to visualize the grouping of genes that overlap in multiple treatments. Treatment intersections are defined by treatments that can be grouped by commonality and, in this context, overlapping differentially expressed genes. Differentially expressed genes (DEGs) are genes that were found to have log_2_FC > 0 and log_2_FC < 0 after a PFAS treatment compared to a control (p_adj_ < 0.05). **B**. Fifteen notable genes which were impacted after exposure to PFOS, PFOA, PFDS, PFDA, 8:2 FTS, and 8:2 FTOH. Of 721 genes that were differently expressed (p_adj <_ 0.05), 10 genes were common to all treatments (red arrow). Five genes that were regulated in at least three treatments had log_2_FC >1. ^*^Genes that, when dysregulated, have been linked to a cancerous pathology. ⸸Genes that, when dysregulated, have been linked to neurodegenerative pathologies. ^**^Genes in this list were impacted in at least three out of six treatments.

Overall, these results highlight several key observations. First, among the PFAS tested, PFOA induced the greatest number of changes in gene expression, suggesting that PFOA is more potent in disrupting cellular homeostasis compared to other PFAS. Second, there does not appear to be a consistent correlation between the level of PFAS uptake (**Figure 1**) and the extent of transcriptomic changes induced by each treatment (**Figure 2-3**). This indicates that it is not the level of PFAS accumulation alone that drives differential gene expression; rather, distinct molecular targets may be involved for different PFAS compounds. Supporting this idea, PFDA and PFOA, both carboxylate PFAS, had the highest overlap in differentially expressed genes, suggesting that the headgroup composition may play a crucial role in determining the cellular effects of PFAS. Finally, our findings identified 11 genes as potential biomarkers for PFAS exposure, as their expression was significantly altered across all six PFAS treatments.

### Potential markers of PFAS exposure

Recent studies have demonstrated that protein expression changes related to neurodegenerative pathologies can be detected in peripheral body fluids, such as blood and saliva.^53, 54^ Furthermore, damage to the BBB can lead to the release of brain-derived proteins into the bloodstream.^55^ Notably, PFAS have been shown to disrupt the BBB, likely through oxidative stress, which may further contribute to the compromise of BBB integrity.^14^

Through transcriptomics analysis, we identified 10 protein-coding genes that were impacted by all six treatments: *Transmembrane protein 106B* (TMEM106B), *transmembrane p24 trafficking protein 7* (TMED7), *T-box 3* (TBX3), *mesencephalic astrocyte derived neurotrophic factor* (MANF), *pyruvate dehydrogenase kinase 1* (PDK1), *BCL2 interacting protein 3* (BNIP3), *isoleucyl-tRNA synthetase 1* (IARS3), *mitotic spindle organizing protein 1* (MZT1), *thioredoxin interacting protein* (TXNIP) and *H4 clustered histone 13* (HIST1H4L). Each of these genes exhibited consistent regulation (either upregulated or downregulated, **Figure 3B**) across all treatments. This uniform response suggests that these genes may serve as promising biomarkers for assessing PFAS exposure, either individually or in combination. Further research is warranted to investigate the regulation of these genes in response to other PFAS variants and to explore their potential as biomarkers of PFAS exposure.

### Potential adverse effects of PFAS exposure assessed by changes in gene expression

We employed multiple strategies to explore the implications of PFAS-induced differential gene expression. To emphasize biological relevance, we considered not just statistical significance but also the effect size of gene expression changes. In addition to the ten genes impacted across all treatments (**Figure 3B**, red arrow), we concentrated on genes that exhibited at least a two-fold change in expression in at least three of the treatments (**Figure 3B**, blue arrow).

Several genes from the resulting list have been associated with various pathologies. These include BNIP3^56^, HIST1H4L^57^, TBX3^58^, MANF^59^ and TXNIP^60^ (observed in all six treatments), early growth response factor 1 (*EGR1*)^61^, solute carrier family 7 member 11 (*SLC7A11*)^58^ and tolloid-like protein 2 (*TLL2*)^62^ (observed in 5 treatments), niban apoptosis regulator 1 (*NIBAN1*)^63^ and fos proto-oncogene (*FOS*)^64^ (observed in 3 treatments). Importantly, MANF and TXNIP have previously been implicated in neuronal health. MANF, which was consistently downregulated in all treatments, is an important protein for survival of neuronal cells, and evidence in rats has suggested that MANF can reverse symptoms of neurodegenerative diseases.^59^ Conversely, TXNIP, upregulated in all treatments, has been linked to oxidative stress, contributing to neuronal death.^60^

### Gene enrichment analysis suggest PFOA can affect neurodevelopment and neurotoxicity

To identify the biological processes impacted by PFAS exposure, we performed gene enrichment analysis. A total of 34 biological processes were significantly enriched (p_adj_ < 0.05, **Table S4**) among the differentially expressed genes. Notably, genes downregulated by PFOA treatment are enriched for neuronal processes, including axonogenesis, regulation of trans-synaptic signaling, modulation of chemical synaptic signaling, and synapse organization (**Figure 4**). This suggest that PFOA may interfere with neuronal networks in SH-SY5Y cells.

**Figure 4.**
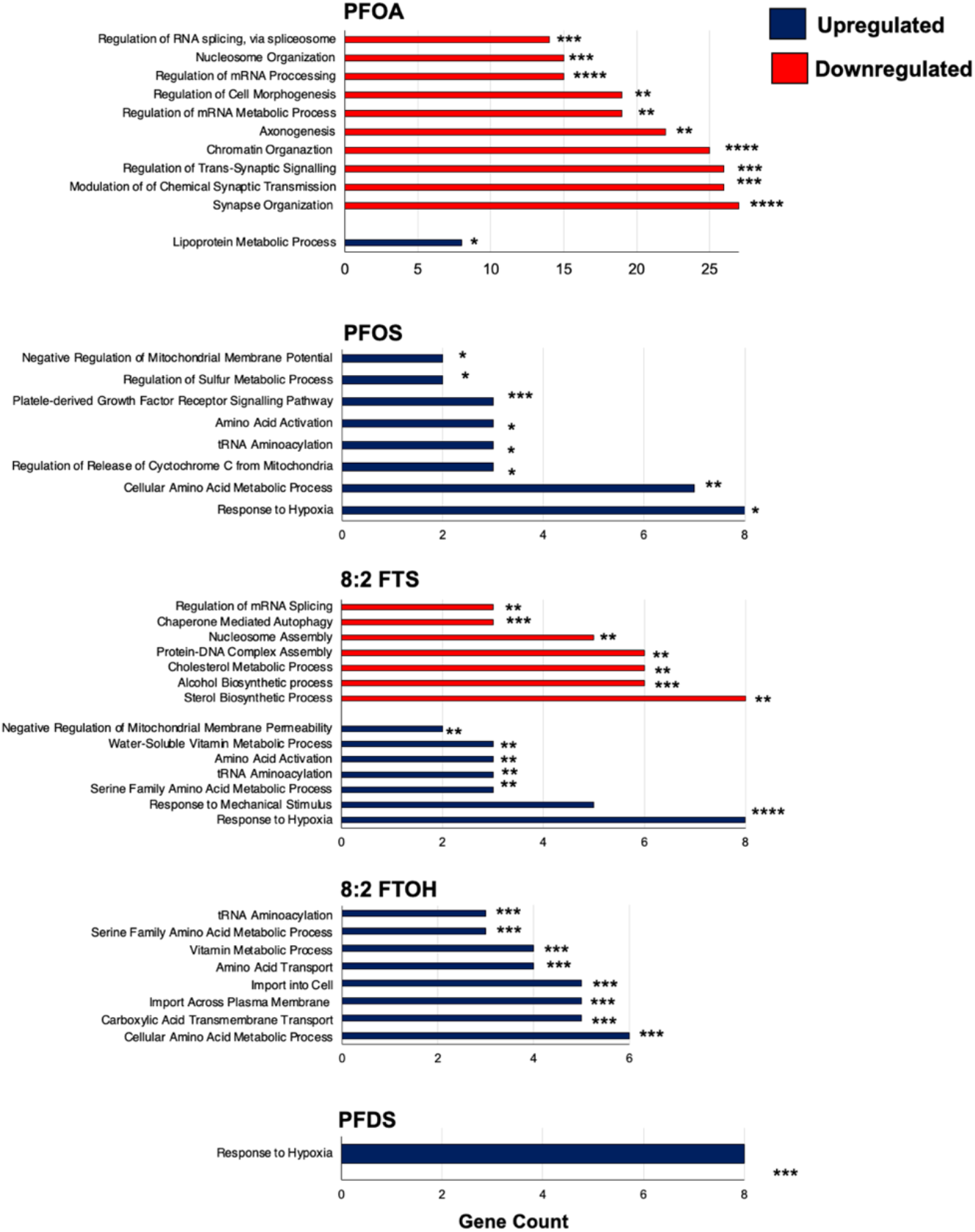
GO gene enrichment analysis for discovering potential biological pathways affected by PFOA, PFOS, PFDS, 8:2 FTS, and 8:2 FTOH treatments. Genes used for this analysis were genes impacted by each treatment with log_2_FC>0.5 (control vs treatment) with p_adj_ < 0.05. The multiple hypothesis corrected p-values are shown in the figure: p_adj_ < 0.05). ^*^p_adj_ < 0.05, ^**^p_adj_ < 0.01, ^***^p_adj_ < 0.001, ^****^p_adj_ < 0.0001. Red and blue indicate enrichments among downregulated and upregulated genes, respectively. Only pathways with p_adj_ < 0.01 are shown (see Table S4 for complete enrichment results).

Consistent with these observations, previous studies have demonstrated that PFOA treatment reduces neurite outgrowth^65, 66^, decreases neurite density and alters nucleolus and mitochondria in neuronal cells^67^. Additionally, PFOA exposure in mice increased expression of brain development-related proteins, and another legacy PFAS, PFOS, was shown to affect learning in zebrafish embryos.^68^ Our findings support the hypothesis that PFAS, particularly PFOA, may disrupt neuronal development and function, with potential implications for neurological disorders.

### Hypoxia- and tRNA aminoacylation-related genes are enriched in differential expression after PFAS treatment

Although the exact mechanisms remain unclear, PFAS exposure can induce a hypoxic state, disrupt cellular energy production, and increase reactive oxygen species (ROS) levels in neuronal cells.^69^ We observed significant enrichment of hypoxia-related genes following treatment of differentiated SH-SY5Y cells with PFOS, PFDS, and 8:2 FTS. These genes include adenylate kinase family (*AK4*), BNIP3, and pyruvate dehydrogenase kinase 1 (*PDK1*), which are linked to hypoxia inducible factor 1 subunit alpha (*HIF-1α*) functioning. Vascular endothelial growth factor A (*VEGFA*)^70^, stanniocalcin-2 (*STC2*)^71^, phorbol-12-myristate-13-acetate-induced protein 1 (*PMAIP1*)^72^, hexokinase 2 (*HK2*)^73^, *SLC2A1*^74^, and DNA damage inducible transcript 4 (*DDIT4*)^75^ with at least one gene being upregulated in each PFAS treatment. These transcriptional changes suggest that sulfonate-containing PFAS, compared to their counterparts with different headgroups, exhibit a higher tendency to induce hypoxia signaling.

Finally, we observed that several genes involved in tRNA aminoacylation, including cysteinyl-tRNA synthetase 1 (*CARS1*), aspartyl-tRNA synthetase 1 (*DARS1*), and *IARS1* were upregulated with PFOS, 8:2 FTOH, and 8:2 FTS treatment. These enzymes facilitate the addition of cysteine, aspartic acid, and isoleucine to tRNA necessary for protein synthesis, and are upregulated in the pro-proliferative state of cancer cells.^76^ Interestingly, PFOS, 8:2 FTOH, and 8:2 FTS treatments also led to the upregulation of amino acid uptake/transport, and metabolism, which are associated with increased cellular energy demands.^77,78^ While further investigations are necessary to elucidate the impact of PFAS on protein synthesis and cellular metabolism fully, these results suggest that PFAS may significantly influence critical cellular processes.

### PFAS treatment impact lipidome of differentiated SH-SY5Y cells

Our results indicate that PFAS induce distinct changes at the transcriptomic level and that different biological processes are enriched based on their chemical structure. To further investigate the effects of PFAS using an additional endpoint assessment, we focused on PFDA, PFDS, 8:2 FTOH, and 8:2 FTS, representing PFAS with different head groups and degrees of fluorination. We then examined the changes these compounds induce in the cellular lipidome. Following a similar protocol as described above, we treated differentiated SH-SY5Y cells with six PFAS at 30 μM and analyzed the alterations in the cellular lipidome compared to the control group (see Methods for sample preparation and lipid analysis details).

We targeted abundant lipid classes, including fatty acids, phosphatidylserines, phosphatidylcholines, diacylglycerols, triacylglycerols, ceramides, and sphingomyelins, with varying acyl chain lengths and degrees of unsaturation. Significant accumulation of fatty acids (∼1.5-2.7-fold, p < 0.05) was observed in cells treated with PFDA, PFDS, and 8:2 FTS (**Figure 5, Table S5**), while phospholipids and diacylglycerols remained unchanged. Treatment with 8:2 FTOH led to modest depletions in several triacylglycerol species (p < 0.05). Sphingolipids also exhibited PFAS-specific changes, with accumulations of certain ceramide species (∼1.3 fold) and sphingomyelins (∼1.6 fold) in PFDA- and 8:2 FTS-treated cells (**Figure 5, Table S5**). Overall, these findings support our conclusions from the transcriptomics analysis, demonstrating that PFAS induce a range of effects at the cellular level depending on their chemical structure.

**Figure 5.**
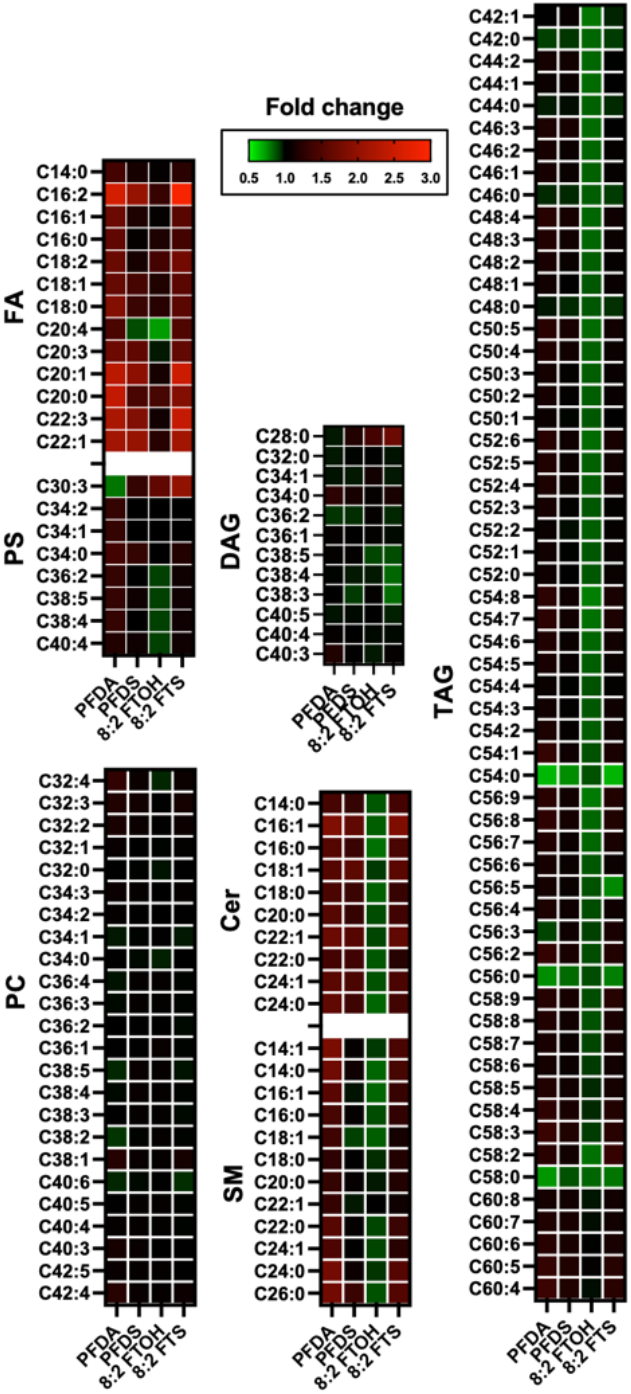
Targeted lipidomics highlight changes in fatty acids and glycerolipids in response to PFAS treatment in differentiated SH-SY5Y cells. Targeted lipids include fatty acids (FA), phosphatidylserine (PS), phosphatidylcholine (PC), diacylglycerols (DAG), triacylglycerols (TAG), ceramides (Cer), and sphingomyelin (SM). The red-to-green heat map shows the fold changes calculated by dividing the abundances of each PFAS treatment by the average abundance of the MeOH control.

## CONCLUSIONS

This study highlights the neurotoxic effects of select PFAS in differentiated SH-SY5Y cells, focusing on complex molecular interactions that may drive PFAS-induced neurotoxicity. The SH-SY5Y model served as a suitable *in vitro* system to investigate the cellular and molecular disruptions caused by PFAS exposure, specifically examining cellular uptake, transcriptomic, and lipidomic alterations.

Our research findings demonstrate that the uptake of PFAS varies significantly among different types. We observed that for both carboxylate and sulfonate head groups, cellular uptake of PFAS increases with the chain length. Notably, PFDS, a fully fluorinated PFAS, shows a more effective uptake than its fluorotelomer counterpart, 8:2 FTS. Additionally, 8:2 FTOH is taken up more efficiently than 8:2 FTS, highlighting the influence of the PFAS head group structure on uptake. We also showed that PFAS induce distinct transcriptomic and lipidomic changes, with varying impacts on neuronal biology depending on their chemical structure. Specifically, PFAS with different head groups and degrees of fluorination modulate various biological processes, including those related to neuronal health and metabolism. Transcriptomic and gene enrichment analyses revealed the enrichment of key pathways involved in hypoxia signaling, oxidative stress, protein synthesis, and amino acid metabolism, all of which are crucial for neuronal function and development. These molecular alterations are consistent with previously observed PFAS-induced impairments in neuronal growth, synaptic signaling, and overall cellular health.

Lipidomic analysis further supported these findings by showing that PFAS exposure leads to specific changes in lipid metabolism, including the accumulation of fatty acids and sphingolipids, both of which are linked to cellular energy homeostasis and membrane integrity. Importantly, these results highlight the diverse cellular responses elicited by PFAS, suggesting that their neurotoxic effects are mediated through complex, multifactorial pathways involving both transcriptomic and metabolic disruptions.

Taken together, our study provides compelling evidence that PFAS can affect neuronal cells at multiple biological levels. Given the widespread use and persistence of PFAS in the environment, further research is critical to fully elucidate their long-term effects on the nervous system. These insights could inform future regulatory decisions and the development of biomarkers for PFAS exposure and toxicity.

## Acknowledgements

We gratefully acknowledge the support from the U.S. Environmental Protection Agency STAR grant (Award No. 84045101). Any opinions, findings, conclusions, or recommendations expressed in this publication are those of the author(s) and do not necessarily reflect the view of the USEPA.

