## Supplemental Information document for "Transcriptomics effects of per- and polyfluorinated alkyl substances in differentiated neuronal cells"

**Table of Contents**

**Table S1:** LC-QToF-MS PFAS analysis parameters

**Table S2:** LC-MS/MS FTOH Analysis parameters

**Table S3:** Genes that showed differential regulation ($p_{adj} < 0.05$)

**Table S4:** GO Gene Enrichment Biological Processes pathways, gene counts, and adjusted p-values

**Table S5:** Targeted lipidomics of PFAS treated differentiated SH-SY5Y cells

***Table S1:*** *Adducts used for the analysis of PFAS.*

| **PFAS** | **Adduct** | ***m/z*** | **PFAS** | **Adduct** | ***m/z*** |
|---|---|---|---|---|---|
| **Native PFAS** | | | **Mass-labelled isotope standard** | | |
| PFOA | $[M\text{-}COOH]^-$ | 368.9765 | M8PFOA | $[M\text{-}COOH]^-$ | 376.0000 |
| PFDA | $[M\text{-}COOH]^-$ | 468.9702 | M6PFDA | $[M\text{-}COOH]^-$ | 473.9869 |
| PFOS | $[M\text{-}H]^-$ | 498.9302 | M8PFOS | $[M\text{-}H]^-$ | 506.957 |
| PFDS | $[M\text{-}H]^-$ | 599.9316 | M8PFOS | $[M\text{-}H]^-$ | 506.957 |
| 8:2 FTS | $[M\text{-}H]^-$ | 526.9615 | M2-8:2 FTS | $[M\text{-}H]^-$ | 528.9682 |
| **Internal Standards** | | | | | |
| M4PFOA | $[M\text{-}COOH]^-$ | 371.9866 | | | |
| M4PFOS | $[M\text{-}COOH]^-$ | 502.9436 | | | |

**Table S2.** Analyte, surrogate standards and transitions and collision energies.

| **Analyte** | **Transitions monitored (*m/z*)** | **Collision Energy (V)** |
|---|---|---|
| 8:2 FTOH | 698.0 → 237.0 | 48 |
| | 698.0 → 252.0 | 33 |
| $^{13}C_2$-6:2 FTOH | 600.0 → 252.0 | 44 |
| | 600.0 → 237.0 | 29 |
| $^{2}H_2,^{13}C_2$-6:2 FTOH | 602.0 → 252.0 | 44 |
| | 602.0 → 237.0 | 29 |

***Table S3:*** *Genes that showed differential regulation ($p_{adj} < 0.05$). This is provided as a separate Excel file.*

***Table S4:*** *GO Gene Enrichment Biological Processes pathways, gene counts, and adjusted p-values*

| | 8:2 FTOH | | |
|---|---|---|---|
| | Biological process | Number of Genes | p-value |
| Up-regulated | Cellular Amino Acid Metabolic Process | 6 | 0.001052048 |
| | Carboxylic Acid Transmembrane Transport | 5 | 0.000571452 |
| | Import Across Plasma Membrane | 5 | 0.001052048 |
| | Import into Cell | 5 | 0.002667565 |
| | Amino Acid Transport | 4 | 0.004472313 |
| | Vitamin Metabolic Process | 4 | 0.002609833 |
| | Serine Family Amino Acid Metabolic Process | 3 | 0.002667565 |
| | tRNA Aminoacylation | 3 | 0.004847855 |

| | PFOA | | |
|---|---|---|---|
| | Biological process | Number of Genes | p-value |
| Up-regulated | Lipoprotein Metabolic Process | 8 | 0.044386031 |
| Down-regulated | Synapse Organization | 27 | 8.14322E-05 |
| | Modulation of of Chemical Synaptic Transmission | 26 | 0.000315295 |
| | Regulation of Trans-Synaptic Signalling | 26 | 0.000315295 |
| | Chromatin Organaztion | 25 | 3.35093E-15 |
| | Axonogenesis | 22 | 0.0030368 |
| | Regulation of mRNA Metabolic Process | 19 | 0.001144803 |
| | Regulation of Cell Morphogenesis | 19 | 0.001953518 |
| | Regulation of mRNA Proccessing | 15 | 2.58602E-11 |
| | Nucleosome Organization | 15 | 0.00045348 |
| | Regulation of RNA splicing, via spliceosome | 14 | 0.000357905 |

| | PFOS | | |
|---|---|---|---|
| | Biological process | Number of Genes | p-value |
| Up-regulated | Response to Hypoxia | 8 | 0.000618741 |
| | Cellular Amino Acid Metabolic Process | 7 | 0.001583171 |
| | Regulation of Release of Cyctochrome C from Mitochondria | 3 | 0.03311181 |

| | | | |
|---|---|---|---|
| | tRNA Aminoacylation | 3 | 0.03311181 |
| | Amino Acid Activation | 3 | 0.03311181 |
| | Platele-derived Growth Factor Receptor Signalling Pathway | 3 | 0.000342707 |
| | Regulation of Sulfur Metabolic Process | 2 | 0.03311181 |
| | Negative Regulation of Mitochondrial Membrane Potential | 2 | 0.045449165 |

| | **PFDS** | | |
|---|---|---|---|
| | Biological process | Number of Genes | p-value |
| Up-regulated | Response to Hypoxia | 8 | 0.001796144 |

| **PFDA** | | |
|---|---|---|
| Biological process | Number of Genes | p-value |

| | **8:2 FTS** | | |
|---|---|---|---|
| Up-regulated | Biological process | Number of Genes | p-value |
| | Response to Hypoxia | 8 | 4.20143E-05 |
| | Response to Mechanical Stimulus | 5 | 0.004169674 |
| | Serine Family Amino Acid Metabolic Process | 3 | 0.005280144 |
| | tRNA Aminoacylation | 3 | 0.010450155 |
| | Amino Acid Activation | 3 | 0.010450155 |
| | Water-Soluble Vitamin Metabolic Process | 3 | 0.011174693 |
| | Negative Regulation of Mitochondrial Membrane Permeability | 2 | 0.011174693 |
| Down-regulated | | Number of Genes | p-value |
| | Sterol Biosynthetic Process | 8 | 4.23644E-07 |
| | Alcohol Biosynthetic process | 6 | 0.000296056 |
| | Cholesterol Metabolic Process | 6 | 0.000315998 |
| | Protein-DNA Complex Assembly | 6 | 0.002113209 |
| | Nucleosome Assembly | 5 | 0.002862356 |
| | Chaperone Mediated Autophagy | 3 | 0.000855188 |
| | Regulation of mRNA Splicing | 3 | 0.002289204 |

**Table S5: Results of lipidomics analysis.** Relative abundances with respect to control group are shown. This is provided as a separate excel sheet.
